# AMP: An Automated Marker Based Motion Capture Processing Pipeline for Kinematic Trajectory Label Scrubbing and Gaps Filling

**DOI:** 10.64898/2026.09.25.754545

**Authors:** Sixu Zhou, Ryan Casey, Joseph Walter, Aaron Young

**Author notes:** Corresponding author *Email address:* (Sixu Zhou).

## Abstract

Marker-based optical motion capture remains the gold standard for kinematic analysis across biomechanical research. However, when tracking dynamic movements, severe occlusions often create missing data and noisy signals that require time-consuming manual cleaning. Existing automated methods usually rely on predefined skeleton models or need massive training datasets for machine learning solutions. This study presents the AMP: an automated marker based motion capture processing pipeline. The AMP is a fully analytical approach that uses hierarchical algorithms including 1) rigid-body constraint, 2) Kalman Filter state estimation and 3) kinematic pattern match to label markers and fill data gaps without manual human intervention. First, the algorithm identifies and removes mislabeling markers before any data scrubbing. During the data cleaning process, our hierarchical algorithm uses a translation-based step to track segments even when there are insufficient donor markers to form a full rigid body. To reconstruct missing marker trajectories, the pipeline initially uses Piecewise Cubic Hermite Interpolating Polynomial (PCHIP) interpolation for short marker gaps. However, the PCHIP creates massive spatial errors during long occlusions, the system automatically switches to rigid-body tracking for gaps over 40 ms. Results showed the pipeline drastically reduced processing time compared to manual human expert methods by 73.5%. In addition, the AMP kept spatial errors within 10 mm of the manual ground truth. By using this combined hierarchical approach to match marker processing accuracy, the pipeline provides a fast, general-purpose tool that fully automates data cleaning and saves hours of manual effort.

## 1. Introduction

Marker-based optical motion capture (MoCap) is a fundamental tool in biomechanics, widely used to investigate human and animal locomotion, build musculoskeletal models, and perform clinical gait analysis (Seth et al., 2018; Henry et al., 2025). During a typical collection session, retro-reflective markers are fixed to anatomical landmarks and tracked by a calibrated cluster of infrared cameras. The resulting 3D marker trajectories are then used to compute inverse kinematics and inverse dynamics from ground reaction force (Delp et al., 2007). Markerless Mo-Cap systems are gaining popularity because they are easier to set up, relying only on a few RGB cameras and computer vision. However, these systems use a pure machine learning approach, indicating their accuracy heavily depends on training data. This data often lacks examples from unique populations or specific activities, limiting how well the systems work across the wide range of conditions that researchers might study (Ceseracciu et al., 2014).

Processing marker-based MoCap data is challenging due to two primary tracking errors: marker mislabeling and trajectory occlusions. Although these artifacts can be systematically classified using derivative analysis and low-pass filtering (Skurowski and Pawlyta, 2022), correcting them still requires a substantial amount of manual post-processing by human experts. This bottleneck becomes more severe in experimental studies including wearable devices. In these scenarios, the hardware frequently occludes anatomical markers and creates extended data gaps. In addition, the reflective components of these devices introduce extraneous infrared noise. This generates “ghost markers” that cause the tracking software to mislabel trajectories by swapping true anatomical trajectories with hardware reflections. Standard commercial software cannot automatically resolve this combination of prolonged occlusions and dense optical noise, requiring researchers to manually reconstruct trajectories frame-by-frame, a time-consuming and burdensome process.

To reduce the burden of manual data processing, early automated approaches attempted to address mislabeling and gapfilling. For the trajectory mislabeling, previous studies typically relied on heuristic thresholds to assign labels to unlabeled point clouds based on expected anatomical distances derived from scaled generalized models (Meyer et al., 2014), while derivative analysis was implemented to detect non-physiological acceleration spikes as an indication of a marker swap (Skurowski and Pawlyta, 2022). In addition, structural constraints were used to predict marker locations to reduce the need for manual interventions (Herda et al., 2001). However, these generalized approaches frequently struggle outside of standard, healthy gait analysis. Complex dynamic movements, severe pathological kinematics, or the introduction fo external hardware require atypical marker configurations that deviate from standard clinical models, reducing the accuracy of generic skeletal fitting. When dense marker clusters exhibit substantial spatial overlap during movements, or when environmental reflections produce “ghost” markers near the true trajectories, simple distance-based heuristics are insufficient for reliable marker discrimination and tracking. Thus, these algorithms may incorrectly label noise or adjacent markers as valid, corrupting the biomechanical analysis.

Once these erroneous labels are scrubbed, the subsequent challenge is gap-filling. Early approaches relied on simple mathematical interpolation, such as linear or cubic splines. These methods showed they maintain acceptable error margins, but only for gaps shorter than 200 ms (Howarth and Callaghan, 2010). For longer occlusions, this interpolation fails to capture rigid-body kinematics, producing peak kinematic errors exceeding 37.1 degrees (Howarth and Callaghan, 2010). To address these larger gaps, previous studies used local coordinate system (LCS) interpolation governed by rigid-body kinematics and reduce the maximum joint angle errors from 23.4 degrees to 9.3 degrees compared to splines methods (Howarth and Callaghan, 2010). Other methods were developed using analytical models such as principal component analysis (PCA) and inter-marker distance correlations (Federolf, 2013; Gløersen and Federolf, 2016). For example, a PCA-based approach reconstructed missing markers with a root mean square error (RMSE) of approximately 5 mm during steady-state walking, even with 50% temporal data loss (Federolf, 2013). While these models effectively utilize kinematic constraints, their performance degrades significantly during macro-occlusions. These algorithms require a minimum number of visible donor markers to track a body segment. When external hardware completely blocks these markers, the localized coordinate system fail, making it impossible to calculate the missing positions (Camargo et al., 2020).

To overcome the limitations of strict analytical constraints, recent research has increasingly turned to data-driven approaches, particularly deep learning and artificial neural networks, to automate marker labeling and resolve trajectory occlusions (Holden, 2018; Kucherenko et al., 2018; Clouthier et al., 2021). These models excel at recognizing complex kinematic patterns and can theoretically reconstruct missing data even when entire marker clusters drop out. However, they function mostly as algorithmic black boxes and require massive, task-specific and patient-specific training datasets. This makes it difficult to apply these models to novel experimental setups, atypical marker configurations, or highly pathological gait patterns that were not heavily represented in the training data. Thus, it suffers from the same set of problems currently experienced by many clinical researchers with markerless Mo-Cap. It is critical to know exactly how a missing trajectory was mathematically reconstructed to ensure the algorithm is not generating non-physiological movement artifacts disguised as valid kinematic data.

Thus, this study introduces a transparent, training-free Mo-Cap processing pipeline adaptable to various datasets. Rather than relying on generalized models or massive datasets, our method uses a subject-specific static calibration to identify and remove erroneous labels. It then resolves trajectory gaps through a hierarchical architecture: rigid-body constraints, Kalman Filter state estimation, and single donor pattern matching. The primary objective was to evaluate the kinematic accuracy and computational efficiency of this pipeline against manual human expert processing. We hypothesize that the automated pipeline would 1) vastly reduce processing time and 2) preserve the kinematics accuracy compared manual human expert methods.

## 2. Methods

### 2.1. Pipeline Architecture Overview

The proposed pipeline processes corrupted MoCap data in two sequential phases to remove optical artifacts from propagating into the final biomechanical analysis (Fig. 1). Following a subject-specific state pose calibration to establish baseline segment geometry, the pipeline first runs a deterministic label-scrubbing phase. This step identifies labeling errors and uses a hierarchical algorithm to correct swapped trajectories or jumps. Lastly, the pipeline transitions to a similar hierarchical marker reconstruction phase to resolve any remaining data gaps.

**Figure 1:**
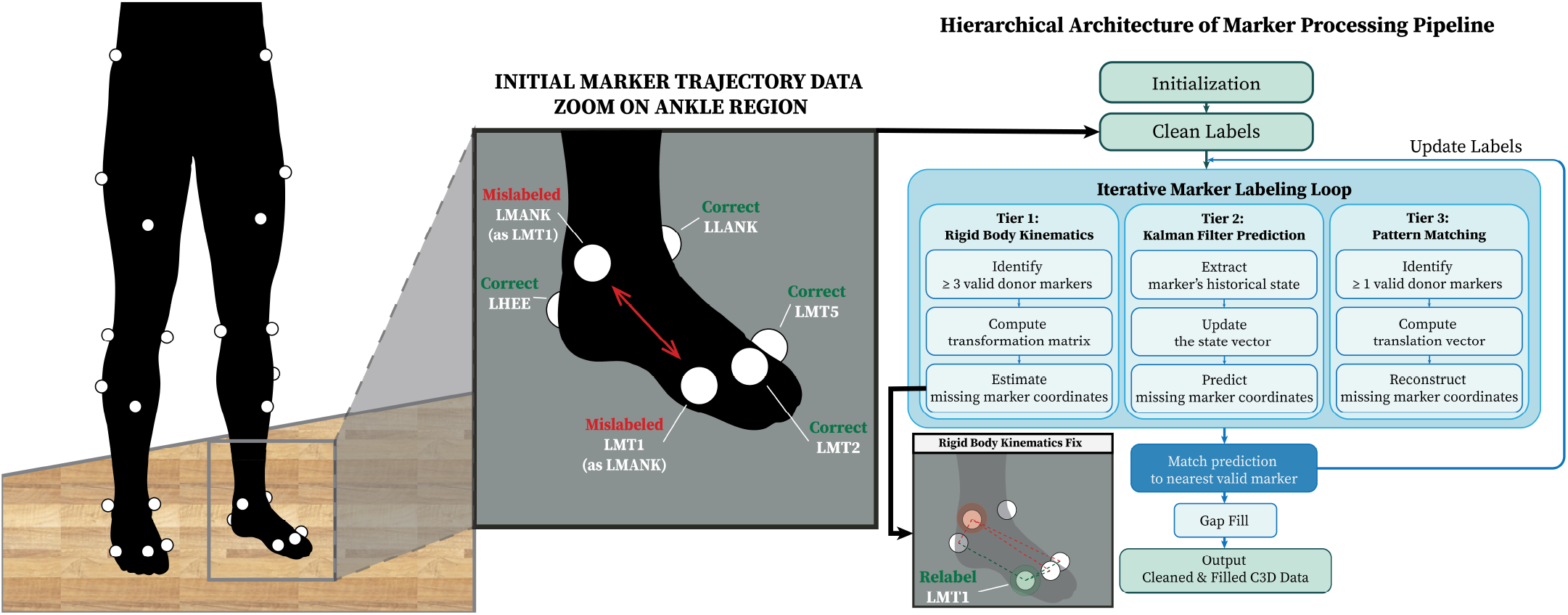
Overview of the proposed hierarchical processing pipeline including the subject-specific static initialization and a three-tiered data-scrubbing architecture.

### 2.2. Static Calibration Reference

Generic skeletal models rely on predefined marker configurations. These templates often fail when markers must be shifted to accommodate physical obstructions or unique subject anthropometry. To accommodate atypical marker configurations, the pipeline directly extracts a custom skeletal template from a subject-specific static calibration trial. For each tracked body segment *j* consisting of *M* markers (where *M* ≥ 3), the algorithm establishes a geometric baseline by calculating the average marker positions from the static calibration trial. Let the averaged global spatial coordinates of the *i*-th marker within the *j*-th segment denoted as **p**_*i, j*_ ∈ ℝ^3^, the exact spatial arrangement of the entire cluster is stored as a constant geometric template matrix:

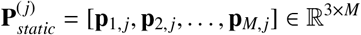

By capturing the exact distances between markers, 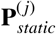 serves as the rigid-body reference for the segment. During dynamic movement trials, the algorithm aligns this rigid template with the moving markers frame-by-frame using a quaternion-based absolute orientation method, enabling the subsequent hierarchical trajectory scrubbing and gap-filling.

### 2.3. Temporal Segmentation and Parallelization

To optimize computational efficiency and leverage modern multicore processors, the dynamic trials are not evaluated as a single, continuous time-series. Prior to the label-scrubbing phase, the pipeline automatically crops the raw MoCap data into smaller, discrete temporal segments. These segments are then distributed and processed simultaneously using parallel computing. The hierarchical prediction and spatial matching algorithms operate independently within each isolated window. Once label assignment and artifact rejection are complete across all parallel threads, the pipeline continuously stitches the segments back together. This multi-threaded architecture drastically reduces the total algorithmic execution time, making the pipeline highly scalable for prolonged dynamic trials.

### 2.4. Hierarchical Label Scrubbing

During dynamic trials, the pipeline generates a spatial prediction, 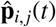, for every expected *i*-th marker on *j*-th body segment at time *t*. Rather than relying on a single gap-filling algorithm, this prediction is generated through a three-tiered hierarchy that adapts instantaneously to the number of valid markers currently visible in the local cluster.

#### 2.4.1. Tier I: Rigid-Body Prediction

When three or more valid markers are present on the *j*-th segment (*d ≥* 3), the system computes the optimal rigid-body transformation between the static geometric template, 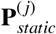, and the corresponding observed dynamic markers using Horn’s closed-form absolute orientation method (Horn, 1987). By representing rotations with unit quaternions, this approach extracts the rotation matrix, 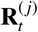, and translation vector, 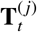. The predicted global coordinate, 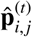, for marker *i* on segment *j* at frame *t* is calculated as in Equation (1):

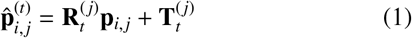

#### 2.4.2. Tier II: Constant Acceleration Kalman Filter

If severe occlusion degrades the cluster to fewer than three valid markers (*d* < 3), rigid-body assumptions fail and the pipeline transitions to a constant-acceleration Kalman Filter (KF). Because rigid-body transformations cannot be computed, this tier estimates the missing marker coordinates based on their individual kinematic state rather than global segment constraints (Aristidou et al., 2008). The KF operates on a constant acceleration model, defining the state vector as 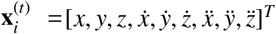. Utilizing the state transition matrix (**F**) and the time step (Δ*t*), the KF propagates the previous kinematic state forward:

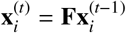

The first three positional elements of the state vector, 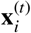, are extracted as the predicted global coordinate, 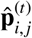.

#### 2.4.3. Tier III: Single-Donor Pattern Match

In extreme cases where the target marker has been occluded too long for the Kalman Filter to maintain a stable covariance, but at least one valid donor marker (*d ≥* 1) remains on the segment, the pipeline switches to a heuristic pattern-matching method. The algorithm computes an instantaneous translation vector (**v**) based on the donor’s displacement over an extended temporal window of *w* frames (where *w ≥* 1). The displacement vector, 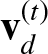, is formally defined as:

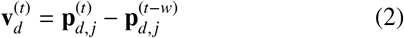

This vector is then applied to the target marker’s last known valid position to generate the final prediction:

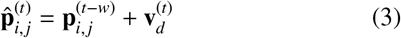

### 2.5. Label Assignment and Error Rejection

Once a spatial prediction 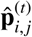 is generated for a target marker, the pipeline evaluates the raw, unlabeled markers for the current frame. To bridge the gap between the mathematical prediction and the physical data, the algorithm employs a nearest-neighbor spatial matching protocol combined with a strict proximity threshold.

Let *U* = {**u**_1_, **u**_2_, …, **u**_*K*_} represent the set of *K* unlabeled markers detected by the optical system at time *t*. The spatial error (the Euclidean distance denoted as *e*_*i,k*_) is computed between the prediction 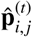 and every available marker **u**_*k*_ ∈ *U* in the unlabeled group:

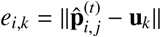

The algorithm identifies the candidate marker **u**_*min*_ that yields the minimum spatial distance. To prevent erroneous assignment to environmental reflections or wearable hardware artifacts, this minimum distance must fall below a predefined proximity threshold, *τ*. This threshold represents the maximum physically plausible displacement of a marker between consecutive frames, governed by the optical system’s sampling frequency and the subject’s baseline kinematics. If *e*_*min*_ ≤ *τ*, the unlabeled marker **u**_*min*_ is assigned the anatomical label of marker *j* and is subsequently removed from the pool of unlabeled groups in *U*. If *e*_*min*_ > τ, the system does not immediately declare the physical marker occluded. Instead, it defers the decision and continues evaluating the rest of the expected markers in the segment. Once all primary assignments are complete, the algorithm performs an additional validation pass on any spatial coordinates remaining in the unlabeled set *U*. If no valid candidate can be identified after all sweeps, or if the unlabeled group is completely empty, the marker is then definitively classified as occluded. This leaves a discrete trajectory gap for that specific frame, which is flagged for final gap-filling in the subsequent reconstruction phase. Any coordinates remaining in the unassigned set *U* are classified as ghost markers. These ghost markers are then ignored from the dataset, ensuring only verified marker coordinates move on to the next processing step.

### 2.6. Hierarchical Gap-Filling Logic

After data scrubbing, the pipeline uses a tiered hierarchy to recover missing marker trajectories, selecting the reconstruction method based on occlusion duration and available segment geometry.

#### 2.6.1. Tier I: PCHIP Interpolation (t_gap_ < 40 ms)

For short-duration gaps, the trajectory of a missing marker 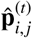 is reconstructed using Piecewise Cubic Hermite Interpolating Polynomial (PCHIP) interpolation, which ensures a smooth transition in both position and velocity at the boundaries of the occlusion. Given the marker coordinates at the beginning (*t*_*i*_) and end (*t*_*i*+*w*_) of the gap, the position during the gap is modeled by a third-order polynomial:

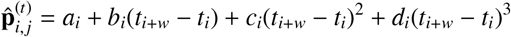

Unlike standard cubic splines that can introduce artificial overshoots within a gap, PCHIP strictly preserves the realistic shape of the trajectory. The coefficients are calculated so the curve matches the actual movement trend at the boundaries.

#### 2.6.2. Tier II: Rigid-Body Constraint Mapping

For gaps longer than 40 ms, the pipeline employs a Tier II rigid-body approach. As long as a minimum of three donor markers remain visible (*d* ≥ 3), the same Horn’s absolute orientation method is used to solve for the segment’s instantaneous rotation 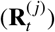 and translation 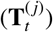. The missing marker *i* is then reconstructed by transforming its static baseline coordinate, **p**_*i, j*_, into the active global space utilizing the transformation established in Equation (1). By shifting to geometric reconstruction for extended durations, the pipeline completely avoids the non-physical artifacts characteristic of prolonged mathematical interpolation.

#### 2.6.3. Tier III: Kinematic Pattern-Fill

When fewer than three donor markers remain visible for a long trajectory gap (*d* < 3), the pipeline utilizes a kinematic pattern-fill. This method assumes the relative spatial displacement between the missing target marker *i* and an adjacent donor marker *d* remains steady over short intervals. To avoid amplifying high-frequency optical noise, the position of the missing marker at frame *t* is estimated by applying the donor’s displacement vector—extracted over a temporal window of *w* frames, as established in Equation (2) and (3)—to the target marker’s last known valid position at the start of that window. This formulation strictly preserves the local kinematic “pattern” of the limb segment.

### 2.7. Experimental Protocol and Ground Truth

To evaluate the efficiency and accuracy of the automated pipeline, experimental data were collected from one able-bodied healthy individual (AB) and one individual with transfemoral amputation (TFAs). All participants provided informed written consent prior to data collection, and the protocol was approved by the Institutional Review Board at the Georgia Institute of Technology. The participants were asked to complete various activities such as treadmill walking, shoveling, cart-pulling, and balance walking. These tasks were selected due to their high dynamic movements. Kinematic data were recorded at 200 Hz using a 38-camera MoCap system (Vicon, UK), and participants were instrumented with a custom retro-reflective marker set consisting of 68 markers. To test the robustness of the pipeline, we used a controlled distortion method. First, we selected high-quality trials with no missing and correctly fully-labeled markers to serve as our ground truth. We then ran a custom script on these trials to intentionally delete marker trajectories and swap labels, mimicking the severe artifacts and errors that happen in a real lab environment. This corrupted data was then fed into the pipeline and compared the repaired trajectories frame-by-frame against the original ground truth to measure the exact error.

### 2.8. Performance Evaluation

To measure the accuracy of the automated pipeline, we directly compared the algorithm’s reconstructed trajectories against the uncorrupted ground-truth data using RMSE of the 3D marker coordinates. For a given marker *i* on segment *j* across a trial of *T* frames, the spatial error between the algorithm’s predicted output, 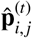, and the true physical coordinate, 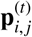, was calculated as:

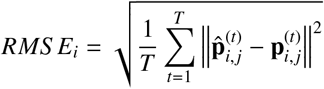

This RMSE was calculated for every marker to determine the overall spatial reconstruction error of the pipeline. In addition to accuracy, we evaluated computational efficiency by recording the algorithm’s total execution time per trial. We then compared this automated processing time directly to the time required by an expert user to manually clean and gap-fill the exact same dataset. The total processing time included the preparation time that requires initial marker reconstructions for all trials.

## 3. Results

### 3.1. Computational Efficiency

The automated script processed a full walking dataset in an average of 37.0 minutes (including 25-minute prep time), whereas an expert researcher required approximately 70.3 minutes (including 25-minute prep time) to manually clean the dataset, showing a 73.5% reduction of time (*p* < 0.05) in data scrubbing process (Fig. 2A). In addition, the manual processing showed a higher accuracy and the AMP achieved under the acceptable 10mm kinematics error (Lin et al., 2020) (Fig. 2B).

**Figure 2:**
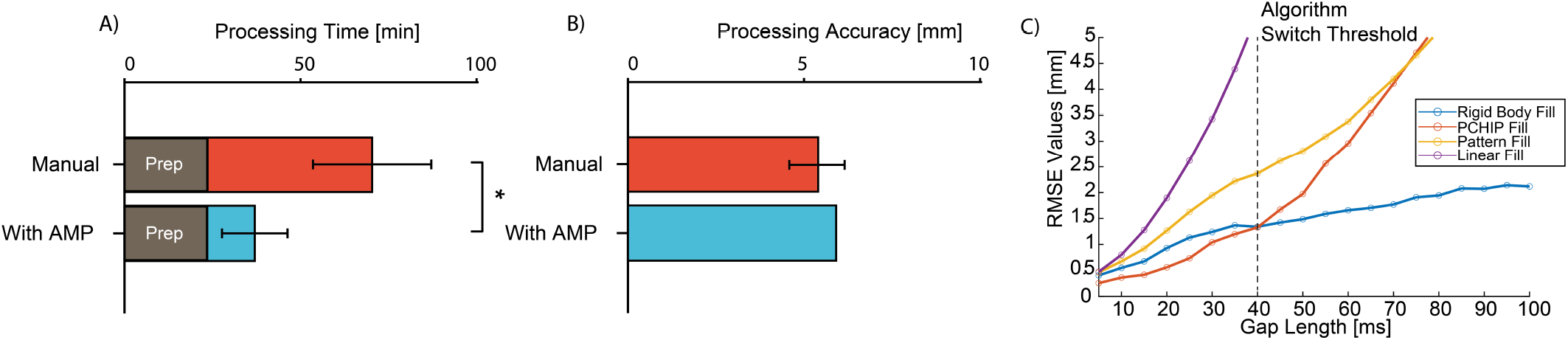
A) Processing Time and B) Processing Accuracy Comparison between Human Experts and AMP Pipeline; C) Algorithm Accuracy vs Missing Gap Length and Algorithm Switch Threshold Determination

### 3.2. Labeling and Reconstruction Accuracy

The pipeline correctly identified and labeled markers in 100% of the frames across all trials. By using a set proximity threshold, the algorithm was able to ignore these “ghost” markers and only assign labels to the actual physical markers. This prevented the label-swapping errors that usually happen when cleaning these types of trials by hand.

Reconstruction accuracy was evaluated across the three tiers of the pipeline, showing a clear performance crossover at the 40 ms threshold (Fig. 2C). For short-duration occlusions under 40 ms, Tier I PCHIP interpolation yielded the lowest spatial error, averaging around 1 mm. However, PCHIP error obviously increased for gaps exceeding 40 ms due to uncertainty and noise of the physical trajectory. For occlusions longer than 40 ms, the Tier II rigid-body method provided superior accuracy by solving for the segment transformation using visible donor markers, keeping error below 5 mm regardless of gap duration. In cases of severe occlusion where fewer than three markers were visible, the Tier III translation-based pattern-fill maintained trajectory continuity. Although this tier produced a higher average error of 10 mm for a gap length up to 100 ms, it served as a necessary failsafe to prevent trajectory termination. The 40 ms search logic was maintained in the final implementation to ensure the pipeline utilized the method with the lowest predicted error for any given gap length.

## 5. Discussion

In this study, the result showed the first hypothesis was accepted — that the automated pipeline drastically reduced processing time by nearly half. Taking advantage of parallel computing techniques, the pipeline indicates the feasibility of reduce manual human efforts. Our second hypothesis was also accepted - that the automated pipeline preserved the kinematic accuracy (*error* ≤ 10*mm*) compared to manual human expert methods. The low average error of marker labeling and reconstruction confirms our hierarchical algorithm is viable in data cleaning without human interventions.

Regarding automated data labeling, our system successfully identified correct markers and ignored real-world reflections without human input. Other studies have attempted to solve this manual labeling problem using deep learning, such as denoising autoencoders (Holden, 2018) or permutation learning (Ghorbani et al., 2019). While these neural networks perform well on standard human walking datasets, they often struggle when researchers must move markers to accommodate atypical marker configurations (Loper et al., 2014; Pan et al., 2024). Neural networks require massive, specific training datasets to learn these new marker layouts (Skurowski and Pawlyta, 2021). Other approaches use LCS to assign labels (Howarth and Callaghan, 2010), similar to our rigid-body approach. However, strict LCS methods fail when a segment has fewer than three visible markers. Our pipeline solves this by adding a translation-based fallback, allowing it to keep tracking and labeling even when a full rigid body cannot be formed. This enables the system to accurately process various marker configurations and complex MoCap data without needing the extensive retraining of neural networks or the perfect visibility required by most methods.

For reconstructing missing data, our pipeline maintained a low average spatial error within 10 mm. Standard MoCap software typically uses cubic spline interpolation to fill gaps (Howarth and Callaghan, 2010). While interpolation works for short missing frames, it creates large spatial errors when a marker is occluded for longer periods (Chiari et al., 2005; Merriaux et al., 2017). To address this, other studies use data-driven methods like PCA (Federolf, 2013) or iterative IK (Camargo et al., 2020). However, IK-based methods adjust marker positions to fit a predefined skeleton model. This process can force sensor noise and skin movement errors (Lavaill et al., 2022) into the final joint angles. Our algorithm avoids this by switching from interpolation to rigid-body tracking for gaps over 40 ms. This maintains high accuracy using only the remaining visible markers, preventing the mathematical overshoots shown in interpolation and avoiding the error propagation common in IK-based methods.

This study has a few limitations. First, the reconstruction method fails when a segment does not have enough visible donor markers for an extended period. Second, the algorithm still requires manual initial labeling at the beginning of the trial to establish the rigid-body templates. Future work will address this by integrating the Hungarian method to automatically assign labels to the initial frames. This update will remove the need for manual setup and make the pipeline fully autonomous.

## 5. Conclusion

Our study investigated the robustness of an automated rigid-body pipeline on processing time and kinematic accuracy when cleaning MoCap data with severe occlusions and artifacts. Our results showed the data scrubbing step successfully removed real-world noise before reconstruction of missing marker trajectories. As part of our hierarchical method, we also used a translation-based step to label markers and track segments even when there are not enough donor markers to form a full rigid body. After scrubbing the data, the reconstruction phase showed that the PCHIP interpolation method had low errors for short missing gaps but generated massive spatial errors during long gaps. The performance crossover at 40 ms shows the need for strict rigid-body tracking. Past machine learning methods required large training datasets and could fail on new marker or skeletal configurations. Our pipeline processed a full dataset within 12 minutes that requires 45 minutes for manual processing and kept spatial errors within 10 mm of the manual ground truth. This pipeline provides a practical and fast tool to automate data cleaning, saving researchers hours of manual work without sacrificing tracking quality.

## Data and Software Availability

The source code for the AMP: Automated Marker Based Motion Capture Processing Pipeline is open-source and available at https://github.com/jasonzhou928/AMP.

## Ethics approval and consent to participate

This study was approved by the Georgia Institute of Technology Institutional Review Board and all participants gave informed consent to participate.

## CRediT authorship contribution statement

**Sixu Zhou:** Software, Formal Analysis, Investigation, Data Curation, Writing – Original Draft, Visualization. **Ryan Casey:** Software, Formal Analysis, Investigation, Data Curation, Writing – Original Draft. **Joseph Walter:** Software, Formal Analysis, Investigation, Data Curation, Writing – Review & Editing. **Aaron Young:** Conceptualization, Methodology, Validation, Writing – Review & Editing, Supervision, Project administration, Funding acquisition.

